# Influenza preimmunity increases vaccination efficacy by influencing antibody longevity, neutralization activity, and epitope specificity

**DOI:** 10.1101/658880

**Authors:** Magen E. Francis, Mara McNeil, Morgan L. King, Ted M. Ross, Alyson A. Kelvin

**Affiliations:** Department of Microbiology and Immunology, Faculty of Medicine, Dalhousie University, Halifax, Nova Scotia; University of Georgia, Center for Vaccines and Immunology, Department of Infectious Diseases, Athens, GA 30602, USA; Department of Pediatrics, Division of Infectious Disease, Faculty of Medicine, Dalhousie University, Halifax, Nova Scotia; Canadian Centre for Vaccinology, IWK Health Centre, Halifax, Nova Scotia

## Abstract

Influenza virus infections are a recurrent public health problem causing millions of hospitalizations each year despite vaccination efforts. The well-known yearly cycling of influenza viruses is the result of the reciprocal and coevolutionary relationship between the host and virus. Together, from the frequent infections and yearly vaccinations humans build a complex immune history over their lifetimes. Despite the prominence of immune history, vaccines are rarely evaluated in the imprinted (preimmune) host. We developed a ferret model for this purpose where ferrets were imprinted with a sublethal dose of the historical seasonal H1N1 strain A/USSR/90/1977 (USSR/77). A +60 day recovery period was given to build immune memory prior to vaccination with a split virion QIV vaccine. To evaluate protection, the ferrets were challenged with a 2009 H1N1 pandemic virus matching the vaccine antigens. The preimmune-vaccinated ferrets did not experience significant disease during challenge while the naïve-vaccinated group were the most severe. Hemagglutination inhibition (HAI) assays showed that preimmune ferrets had a faster and longer antibody response post vaccination for all vaccine antigens compared to minimal HAI responses in the naïve-vaccinated group. To investigate the immune mechanisms leading to disease protection in the preimmune ferrets, we performed microneutralization and isotype ELISA assays. Microneutralization suggested preimmune ferrets developed antibodies that were more functional for virus neutralization. Antibody isotype profiling indicated that virus specific antibodies in the preimmune-vaccinated ferrets was dominated by the IgG isotype suggesting B cell maturity and possible plasticity in a pre-existing B cell. Surprisingly, the naïve-vaccinated ferrets developed a more severe disease with less virus neutralization suggesting improper immunological processing of vaccine antigens. Together, these results showed the preimmune host had greater responses to vaccination, and the predominant IgG virus specific antibodies suggested a flexible long-lived B cell response. These results are important and should be considered for vaccine design.

**AUTHOR SUMMARY:** The influenza virus is a significant threat to human health and the economy despite large-scale vaccination efforts. The low effectiveness of the seasonal influenza vaccine is attributed to the frequently mutating virus enabling people to have several influenza virus infections throughout their lifetimes. As people are susceptible to multiple infections, they build a complex immune history. Despite this, vaccines are often not evaluated in animals with an immune history. Here we developed a ferret model that had previously been infected with a historical influenza virus to evaluate vaccine responses to current vaccines. Ferrets were infected with a sublethal does of a historical virus, A/USSR/90/1977, to develop a preimmune background. Preimmune ferrets were vaccinated with the Sanofi quadrivalent influenza vaccine and the antibody responses were investigated after vaccination. Our results showed that preimmune ferrets had a stronger antibody response following vaccination and the antibodies developed were older and better at neutralizing influenza virus at a virus challenge. Clinically, preimmune-vaccinated ferrets developed a milder disease during challenge compared to naïve-vaccinated ferrets. This work indicates that the host responses to vaccination are dependent on the host background and that influenza vaccine development and evaluation should take host influenza background into account.

## INTRODUCTION

Influenza A virus infection is a recurrent and unsolved public health problem. Two types of Influenza viruses (Orthomyxoviridae), influenza A and influenza B, each with its own subtypes and lineages, currently circulate in humans [1–3]. These viruses cause millions of hospitalizations and thousands of deaths yearly infecting between 5% and 30% of the global population [4–8]. Furthermore, infants, the elderly, pregnant women, and people with pre-existing medical conditions are at higher risk for developing severe disease requiring hospitalization from influenza infection [9].

The well-known yearly cycling of the influenza virus is the result of the reciprocal relationship between the host and virus: the human immune responses influence virus mutation which feeds back to influence immune response [8]. As a result, vaccines need to be reformulated every year, at great expense, to match circulating strains [10]. The changes in the influenza virus occurs through a process known as antigenic drift [11]. Due to antigenic drift, distinct seasonal strains emerge each year with the ability to infect a vulnerable human population [12]. Humans build a complex immune history over their lifetimes as they are cyclically infected with novel strains and receive seasonal vaccinations. It is recognized that immune history or *preimmunity* significantly influences vaccine and infection outcomes, but the mechanisms that regulate vaccine responses in the preimmune host have yet to be elucidated [13, 14].

The host’s primary infection with an influenza virus initiates a cascade of innate and adaptive immune events that culminates in immunological memory. This first infection in a person’s lifetime is referred to as the viral imprinting event [15]. The goal of the immune response is to mobilize adaptive immunity for the production of antibodies capable of virus neutralization. This response is defined by the generation of short-lived plasma cells (antibody producing cells) [16]. Activated B cells mature into plasmablasts and plasma cells, producing antibodies targeted at viral epitopes. The adaptive cellular and humoral responses move through three phases: expansion, contraction, and memory. The expansion phase is the proliferation of antigen-specific T and B cells producing a large number of reactive cells to control the pathogen. Experimental studies from our group as well as others investigating immune responses in animal models (NHP and ferret) have shown that lymphocytes as well as virus-specific antibodies circulate at high numbers (in the expansion phase) even after pathogen clearance (up to 48 days) [17, 18]. Eventually, adaptive immune cells contract until there is a limited number of highly specific cells able to circulate for immune surveillance, which is referred to as the B cell memory pool [19]. Importantly, the memory phase is the time period in which humans are typically re-exposed either through reinfection or vaccination.

Immune history has an impact on host influenza vaccination responses [15, 20, 21]. These human studies, based on serology or epidemiological study designs, suggest that understanding how the host interacts with an antigenically divergent pathogen, such as influenza viruses over multiple exposures will be important for identifying susceptible populations and designing the next generation of influenza vaccines. Here we investigated the responses to the split, inactivated virion Sanofi QIV influenza virus vaccine (Sanofi Pasteur, Canada) in preimmune ferrets. We first established preimmunity over 67 days to ensure the ferrets were outside of the expansion phase of the adaptive immune response. Vaccinated ferrets that were previously infected with the historical seasonal influenza virus strain A/USSR/90/1977 had significantly divergent responses in terms of antibody reactions and clinical disease compared to naïve ferrets - vaccinated ferrets. These results have implications on future vaccine design and evaluation.

## RESULTS

### Study Design

With the exception of neonates and young infants, the majority of the human population has been previously exposed to influenza viruses. By living through the winter influenza season or any time year-round in the tropics, people will be exposed to influenza viruses either from community transmission or vaccination; therefore, understanding vaccine responses in the preimmune host will give a more accurate insight into the responses to vaccination for the majority of the population. The early stages of influenza infection are defined by the antiviral response and innate immune cell activity [22]. Subsequently, adaptive cellular and humoral responses move through phases of expansion, contraction, and memory, to establish host long-term protection (**Figure 1A**) [19]. To understand how the preimmune host responds to vaccination, we designed a study using the ferret model to build an influenza preimmune background for subsequent vaccination and challenge. The design and experimental timeline of the study are shown in **Figure 1A**. The key to our study design was the elongated recovery period. Previous studies investigating the immune re-encounter with influenza viral antigens or live virus infection are frequently designed with short recoveries of 14 to 28 days separating primary and secondary challenges [23–25]. These time frames do not properly account for the actual immunological recovery. Our rationale for this study was to allow the ferrets to recover over several months. Several studies have shown that influenza virus-specific adaptive immune cells exist in high numbers in circulation even after 21 days post infection [17, 18]. Our design includes a long recovery period so that re-exposure (infection or vaccination) occurs after the contraction phase to more accurately reflect the seasonal exposures of influenza viruses in humans.

**Figure 1.**
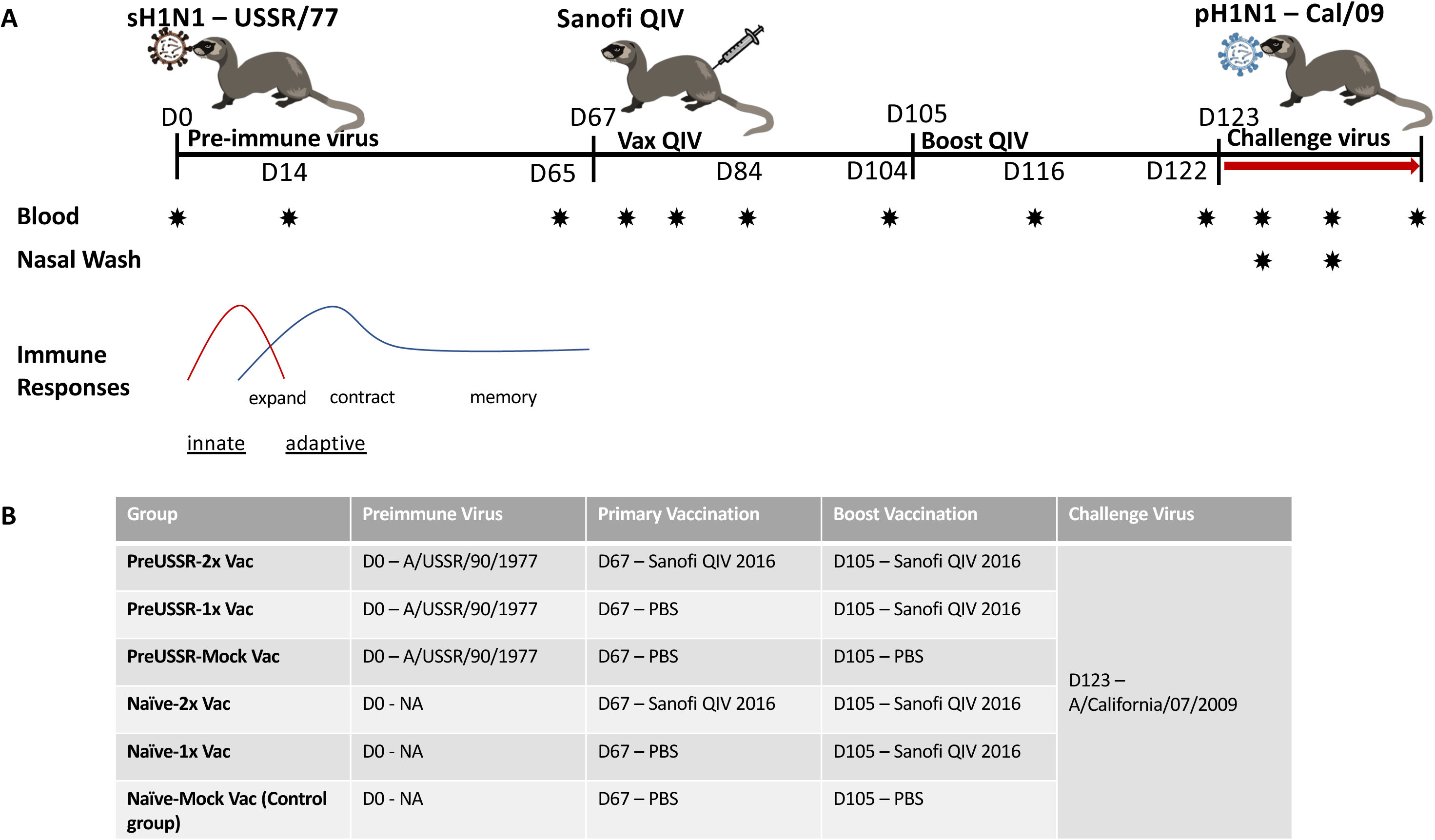
Study timeline and experimental group design for the investigation of the influence of influenza immune history on vaccines responses to Sanofi FLUZONE^®^ QIV.

To establish preimmunity, adult ferrets were infected (imprinted) with a sublethal dose of the historical seasonal H1N1 A/USSR/90/1977 (USSR/77) Day 0 of the study. USSR/77 was chosen as the imprinting virus since it was a previously circulating human influenza virus that would have significantly impacted the immune history of many people alive today [26]. Furthermore, USSR/77 is a virus that re-emerged in the 1970s and is antigenically divergent compared to our proposed contemporary challenge virus [14]. Ferrets recovered over 67 days to decrease non-specific immune responses. The ferrets were subsequently vaccinated at Day 67 post-imprinting infection (pi) and boosted on Day 105 pi with the Sanofi QIV split virion vaccine (FLUZONE^®^ Sanofi-Pasteur, PA, USA). To control the experiment, 6 groups (6 ferrets per group) were designed: **1.)** preimmune USSR/77 – QIV vaccinated and boosted (preUSSR-Vac2x); **2.)** preimmune USSR/77 – QIV vaccinated with no boost (preUSSR-Vac1x); **3.)** preimmune USSR/77 – mock vaccinated (preUSSR-Mock Vac); **4.)** naïve – QIV vaccinated and boosted (Naïve – Vac2x); **5.)** naïve – QIV vaccinated with no boost (Naïve – Vac1x); and **6.)** naïve - mock vaccinated (Naïve – Mock Vac) (**Figure 1B**). To investigate protection from vaccination or preimmune infection, ferrets were infected with a currently circulating 2009 H1N1 pandemic virus on D123 pi. This virus, A/California/07/2009 (Cal/2009), is one of the components of the Sanofi^®^ QIV vaccine used in our study. Furthermore, this strain is a representative from the 2009 H1N1 influenza pandemic which marked the emergence of an antigenically novel H1N1 “lineage” [27]. Since the vaccine contained Cal/2009 antigen, vaccinated animals should be protected against challenge with Cal/2009. Considering the USSR/77 infection and Cal/09 reinfection, this combination represents a monosubtypic heterologous reinfection. Importantly, these viruses do not produce cross-reacting HAI antibodies when infected into the naïve ferret model as we have previously shown [26]. The protein alignment in the highly variable region of HA (135-295aa), which belongs to the HA-RBD area, has only 59% homology between Cal/09 and USSR/77 [26] and this combination of preimmune exposure and challenge virus infection represents conditions plausible for the immune history of people alive today. Blood (serum), lungs, and nasal wash were collected at specified time points to examine the immune response. Specifically, blood samples were collected at Days 0, 14, 65, 68, 72, 79, 104, 116, 122 pi and pc at Days 2, 7, and 14. These time points were chosen as they represented significant days surrounding the infection and vaccination events, as we have previously shown [28, 29].

### Milder clinical disease was observed in the preimmune-vaccinated ferrets at challenge

Three groups of ferrets were infected with 1×10^6^ EID_50_ of the historical H1N1 strain USSR/77 at the start of the study and the temperature and weights were monitored in the ferrets over 14 days post imprinting (pi) (**Supplemental Figure 1**). The remaining ferrets in other groups were left in control rooms to be age matched for subsequent time points in the study. Analysis of the temperature and weight change of each group indicated minimal variation between groups following imprinting infection. All groups lost a moderate amount of weight following infection. At nadir, all groups had lost between 5 and 10% of their original weight similar to our previous findings with this virus [26]. All groups had a temperature increase peaking Day 2 pi. On Day 67 and Day 105, the ferrets were vaccinated and boosted, respectively, as specified by their group (**Supplemental Figure 2**). Minimal weight loss or temperature change was observed over a 14-day monitoring period post vaccination and boost similar to monitor for reactogenicity.

To determine how well the vaccine was able to protect against disease at challenge, on Day 123 the ferrets were challenged with an infectious dose of 10^6^ EID_50_ of A/Cal intranasally. The ferrets were monitored for signs of clinical disease including weight loss, fever, and lethargy post challenge (pc). The preUSSR-Vac2x group lost minimal weight. Maximum weight loss was observed on Day 2 pc where weight dropped to 97% of original weight (**Figure 2A**). Furthermore, the preUSSR-Vac2x did not have a temperature increase pc, but instead experienced a slight drop in body temperature (**Figure 2B**). Mild disease was also experienced by the preUSSR-Vac1x group although the time period at which this group had weight loss was longer than that seen in the preUSSR-Vac2x group and maximum weight loss was noted at 96% of original weight. No temperature increase was observed in the preUSSR-Vac1x group. Conversely, the naïve-Vac2x and naïve-Vac-1x groups both experienced significant temperature increase to 105% and 103% of original temperature at Day 2 pc, respectively (p<0.05 by t-test compared to preUSSR-Vac2x). These groups, as well as the control group (naïve-MockVac), had significant weight loss compared to the preUSSR-Vac2x group. Interestingly, the naïve-Vac2x group had the largest amount of weight loss pc where maximum weight loss was seen on Day 9 pc at 82% of original weight. Together, these results suggest that preimmune ferrets had a greater vaccination efficacy as shown by a milder clinical disease.

**Figure 2.**
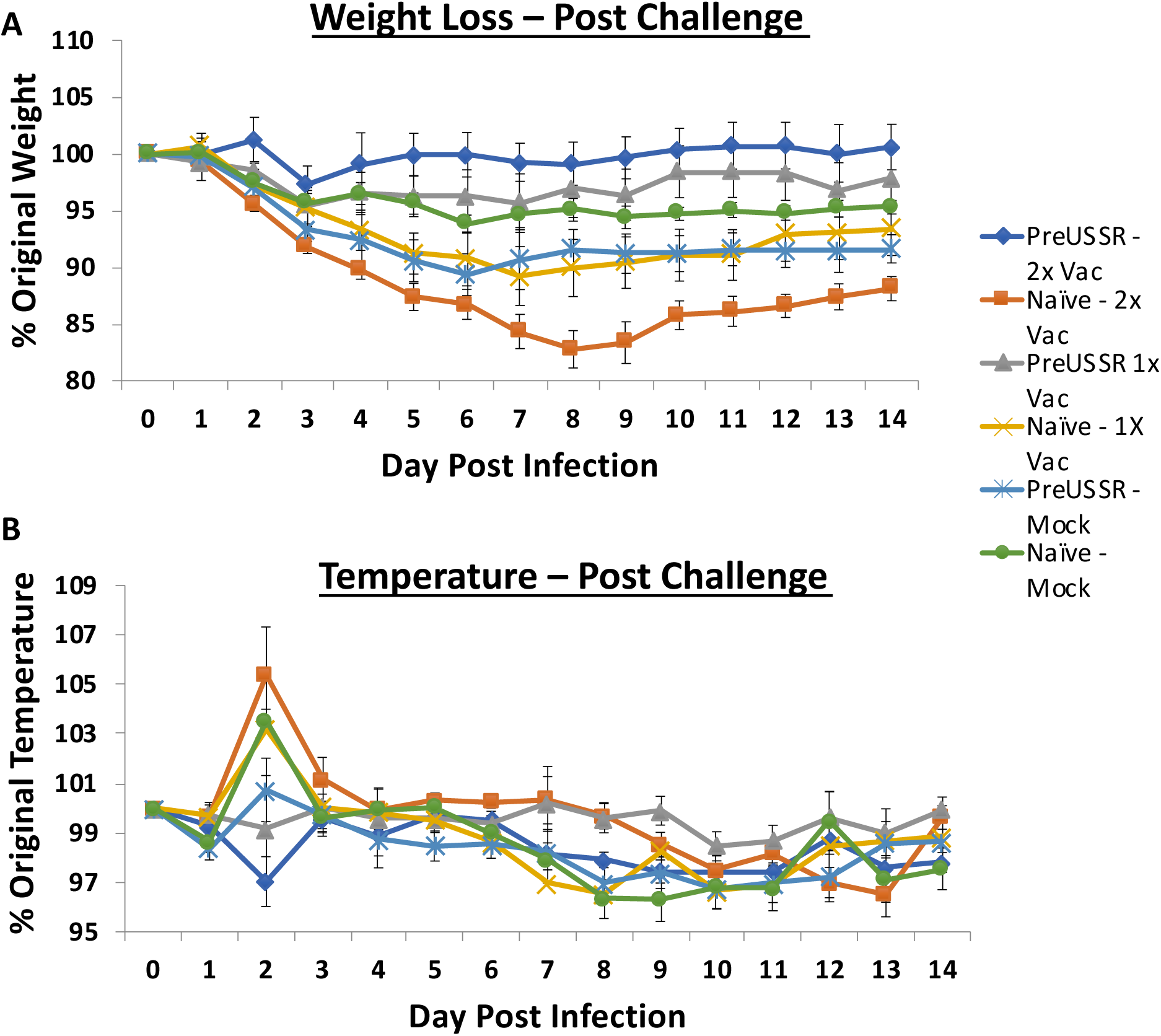
Prior exposure to a heterologous H1N1 influenza virus increases influenza vaccine protection. Ferrets were infected with USSR/77 (10^6^ EID_50_) to establish an influenza-specific immune background or were left naïve. After 67 and 105 days, select groups of ferrets were vaccinated with the Sanofi FLUZONE^®^ QIV vaccine and challenged at Day 123 with the Cal/07 H1N1 pandemic 2009 virus. Weight loss (upper panel) and temperature changes (lower panel) were monitored for 14 days pc. * indicates a p-value of less than 0.05

### Decreased respiratory infection in preimmune-vaccinated ferrets

Hallmarks of influenza virus induced illness include respiratory virus shedding and lung pathology. To determine the viral susceptibility and respiratory disease, we collected nasal washes (NW) and lungs from ferrets pc with the Cal/09 H1N1 pandemic virus. NW were collected Days 2 and 7 pc. Virus was only detected in NW collected on Day 2 pc and is shown in **Figure 3**. Titers in the pre-USSR-vaccinated groups were markedly or significantly lower than the naïve-vaccinated or naïve-unvaccinated control groups. Both preUSSR-vaccinated groups had viral titers averaging at 2.5 log_10_ TCID_50_. Furthermore, the naïve-Vac2x group had the highest titer of viral shedding (**Figure 3**). Histopathological analysis of the lungs collected on Day 14 pc with Cal/09 suggested distinct respiratory responses dependent on influenza immune background. Lungs were collected at necropsy and prepared for histopathological analysis (H&E staining). Stained and mounted lung sections were viewed at High (20x) and Low (5x) Magnification (**Figure 4**). The Naïve-MockVac group showed typical Day 14 pathological features of Cal/09 infection including mononuclear cell infiltration, bronchiolar wall thickening, and hemorrhage (left panels)[30]. The preUSSR-MockVac lungs had evidence of necrotising alveolitis (middle left panels). The preUSSR-Vac2x group ferret lungs had evidence of either mild pathological changes with minimal leukocyte infiltration, peribronchiolar thickening, and alveolar wall thickening (mild) or significant pathological features of interstitial pneumonia with alveolar wall thickening, edema, and fibrosis (affected) (right panels). Taken together, the analysis of viral load shedding and histopathology suggested that the outcomes in the respiratory tract were dependent on the immune background of combinations of vaccination and pre-infection.

**Figure 3.**
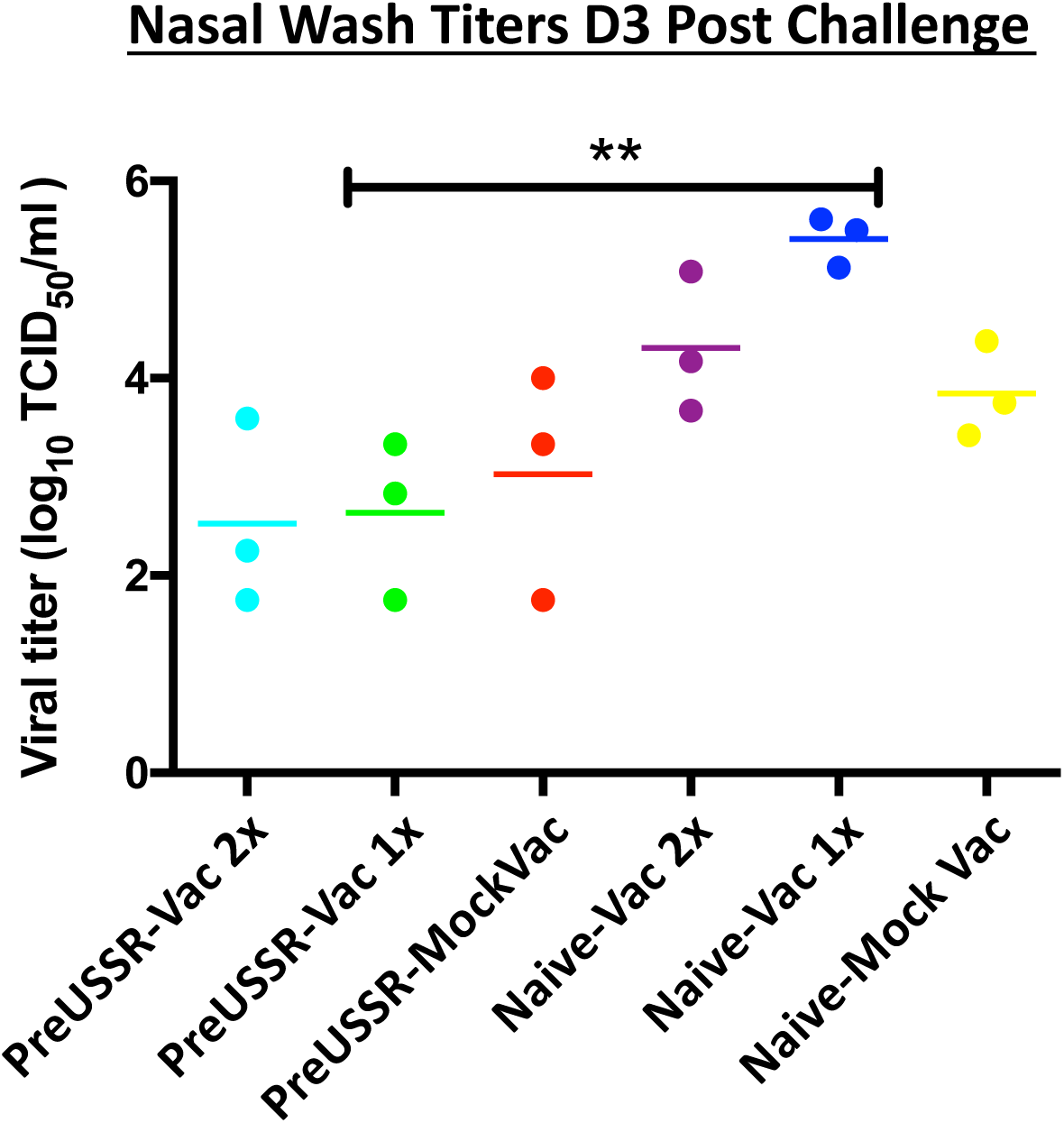
Viral shedding was lower in previously infected and vaccinated hosts. Nasal washes were collected from all groups post Cal/09 challenge on Day 2 pc. Live viral load was calculated by TCID_50_ titration assay using MDCK cells. Student’s t-tests were performed among groups to determine statistical significance. ** indicates a p-value of less than 0.001 between preUSSR-Vac1x and naïve-Vac1x

**Figure 4.**
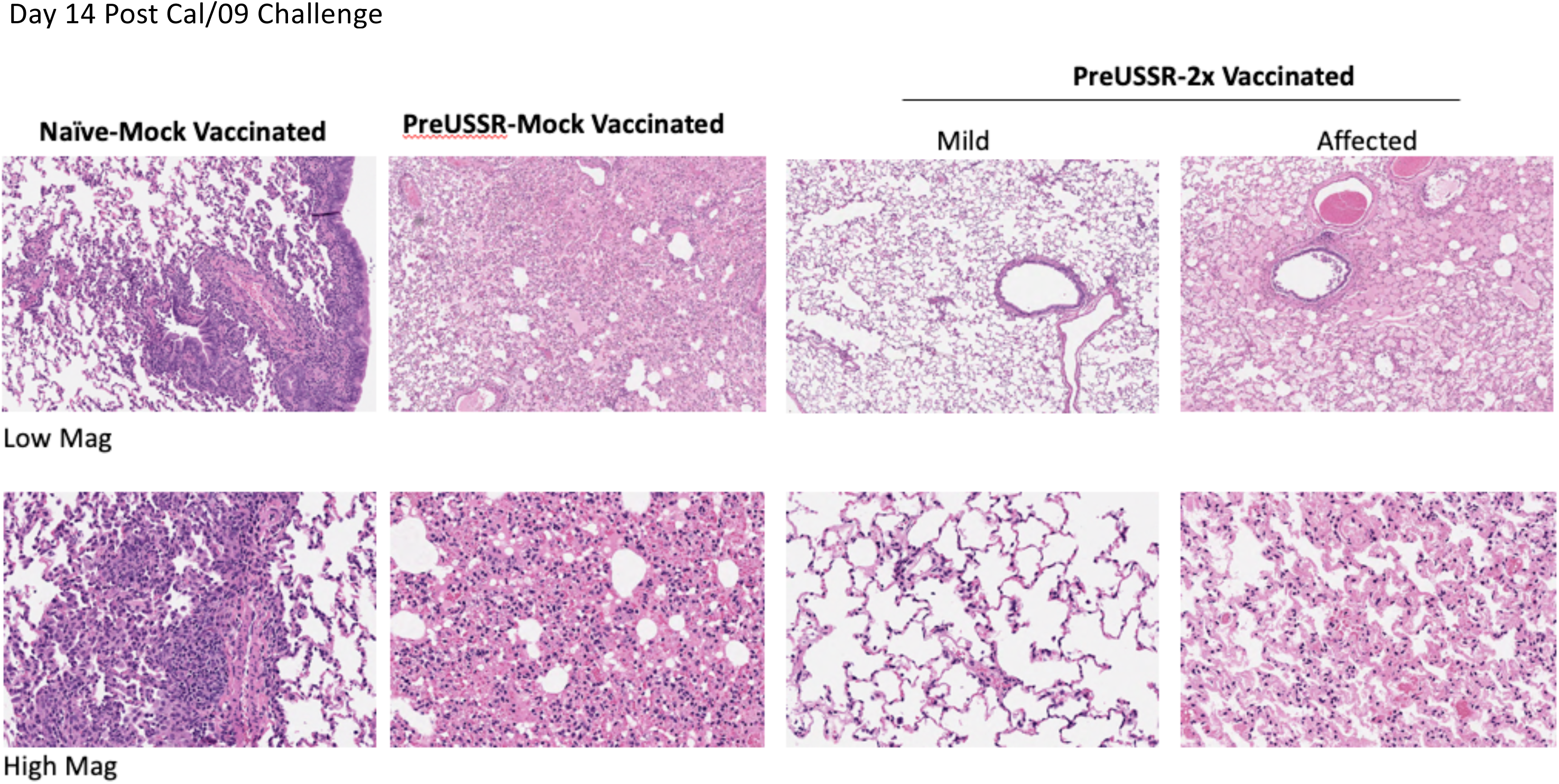
Histopathological analysis shows differential lung pathology dependent on pathogen and vaccination history. Harvested lungs from all ferrets were processed for histopathological assessment. Tissue morphology was assessed by hematoxylin & eosin staining. Data was collected from at least three ferrets per group and results are a representative of the inoculations/infections. High resolution scans were performed using an Aperio ScanScope XT (Leica Biosystems) at 40x magnification. Images were captured using the HALO program from UHN AOMF (Advanced Optical Microscopy Facility) at 5x (low) or 20x (high) magnification of the scan.

### Humoral immunity is differentially regulated in the preimmune host

Serum antibody titers reactive toward the HA protein are often indicative of how protected an individual will be toward a specific influenza virus strain. We next investigated the HA specific antibody responses in each experimental group over the experimental time course to determine if influenza background influences the antibodies elicited following vaccination. Sera was collected from the experimental groups along the experimental timeline on Days 0, 14, 65, 84, 104, 122, 125, 130, 137. Standard HAI assays were performed to quantify antibody specificity and regulation against influenza HA antigens (**Figure 5** and **6**). USSR/77 virus was used to quantify antibodies reactive to this antigen, whereas the 2016 WHO kit was used to quantify antibodies toward the vaccine antigens: 2009 H1, H3, B-Yamagata, and B-Victoria antigens.

**Figure 5.**
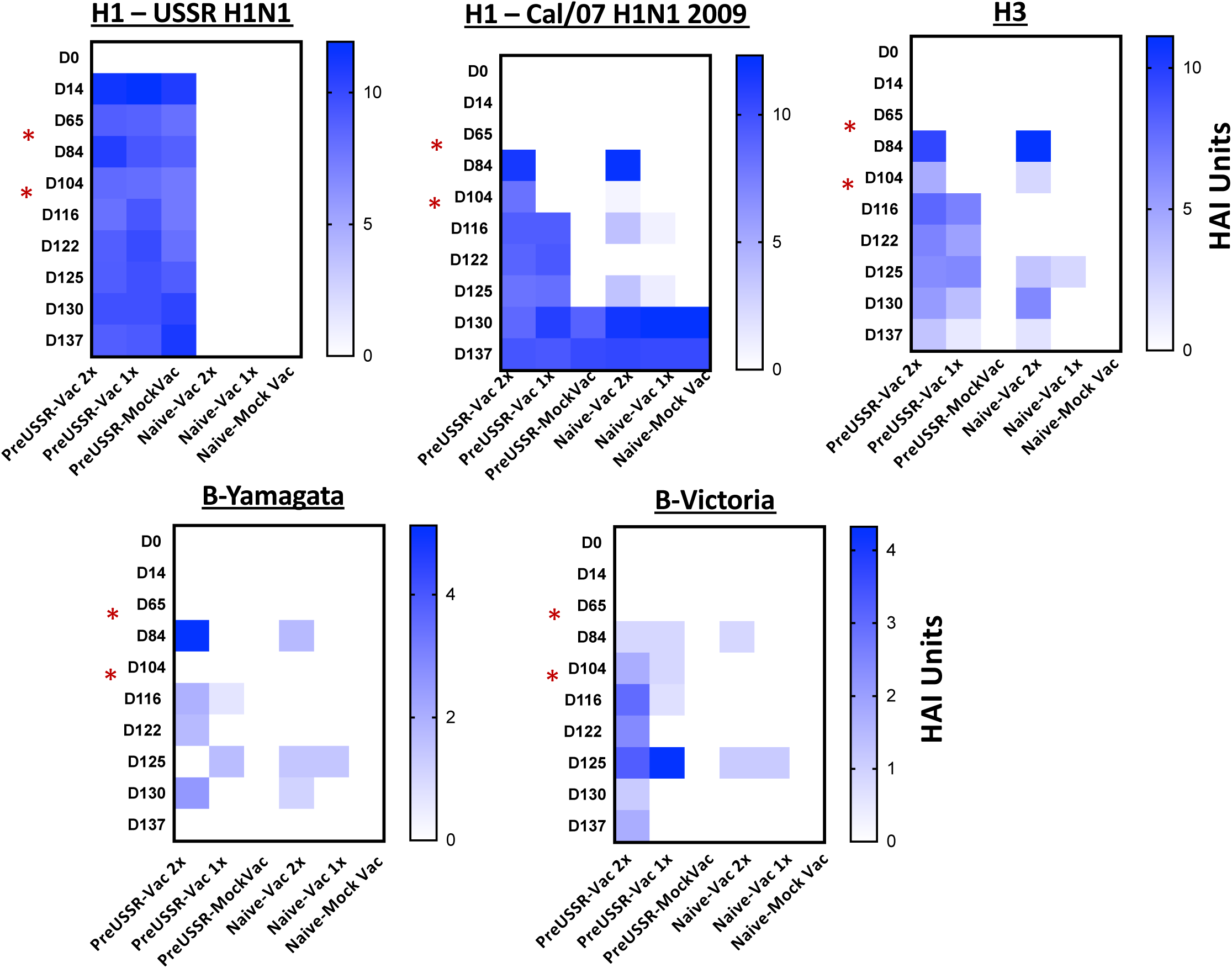
Increased vaccine antibody titer in previously infected ferrets. The mean HAI U for each experimental group was calculated and the results were plotted in heat maps to visually represent changes in antibody titer and specificity over time. HAI assays were performed using specific whole virus (USSR/77) or BPL inactivated vaccine antigens and turkey red blood cells. Previously infected ferrets produced antibody responses of greater titer and longer lived in circulation compared to naïve-vaccinated ferrets. Red arrow indicates vaccination days.

Heat maps of the calculated HAI units were generated to visually summarize the HAI data against seasonal USSR H1, pandemic 2009 H1, H3, B-Yamagata, and B-Victoria HA antigens over the 137-day time course (**Figure 5**). The mean HAI units per group per time point was calculated over the entire study. The H1-USSR heat map (top left graph) shows strong HA antibody generation by Day 14 following infection for all infected groups. All ferrets infected with USSR/77 (H1-USSR) developed HA binding antibodies to the H1-USSR antigen between 10 and 12 HAI Units as assessed on Day 14 pi. Titers in these previously infected animals remained between 7 and 12 HAI Units for the entire study. The uninfected ferrets remained negative throughout the study. At vaccination and boost (Day 67 and Day 105, respectively), the USSR infected ferrets had strong HA antibody responses shown by high HAI titers to the H1 2009 HA protein (top middle graph). These H1 2009 antibody titers generated following vaccination in the preUSSR group remained high throughout the time course. Following first vaccination, both previously infected and naïve ferrets developed high antibody tiers to the H1 2009 HA (H1-2009), between 10 and 14 HAI Units, shortly after vaccination (Day 14). When titers were quantified in sera collected at a longer time point (30 days) following vaccination, the preimmune ferrets had sustained antibody values between 6 and 10 HAI Units whereas 4/5 of the naïve-vaccinated ferrets had undetectable levels of H1-2009 reactive antibodies. Fourteen days following boost, the preUSSR-Vac2x ferrets had H1-2009 titers between 8 and 10 compared to lower values seen in the naïve-Vac2x ferrets (between 3 and 4 HAI Units). Furthermore, H1-2009 antibody titers assessed prior to vaccination showed no detectable levels and between 8 and 11 for the naïve-Vac2x and preUSSR-Vac2x ferrets, respectively, suggesting no cross-reactivity was detected by HAI for antigens Cal/09 and USSR/77. The naïve-Vac1x ferrets also had strong or moderate responses to the vaccine directly following vaccination but these titers quickly waned and were not maintained over time. Interestingly, these trends were consistent for the other HA antigens H3, B-Yamagata, and B-Victoria, where the preUSSR ferrets had a stronger antibody response following vaccination and boost despite the antigenic divergence of the vaccine antigens with the original imprinting virus. Taken together, these results suggest that a previous infection to the influenza viruses primes the naïve immune system to have greater antibody responses following vaccination with the Sanofi QIV^®^ vaccine administered by intramuscular injection.

To visualize numerical changes in antibody titers across the experiment for the H1 2009 antigen, HAI values were determined for each ferret and the group averages were calculated per time point and plotted in a histogram focused on only the preUSSR-Vac2x and the naïve-Vac2x groups (**Figure 6**). Serum collected on Day 84 following first vaccination showed similar HAI levels between the two groups (>11 HAI Units). Interestingly, the subsequent blood collection prior to boost indicated a significant drop in 2009 H1 antibody titer in the naïve-Vac2x groups to <2 HAI Units whereas the PreUSSR-Vac2x group had a minimal drop in HAI to 8 HAI Units. Statistical differences in HAI titers for the 2009 H1 antigen between the preUSSR-Vac2x group and the naïve-Vac2x group were seen throughout the time course until Day 7 pc.

**Figure 6.**
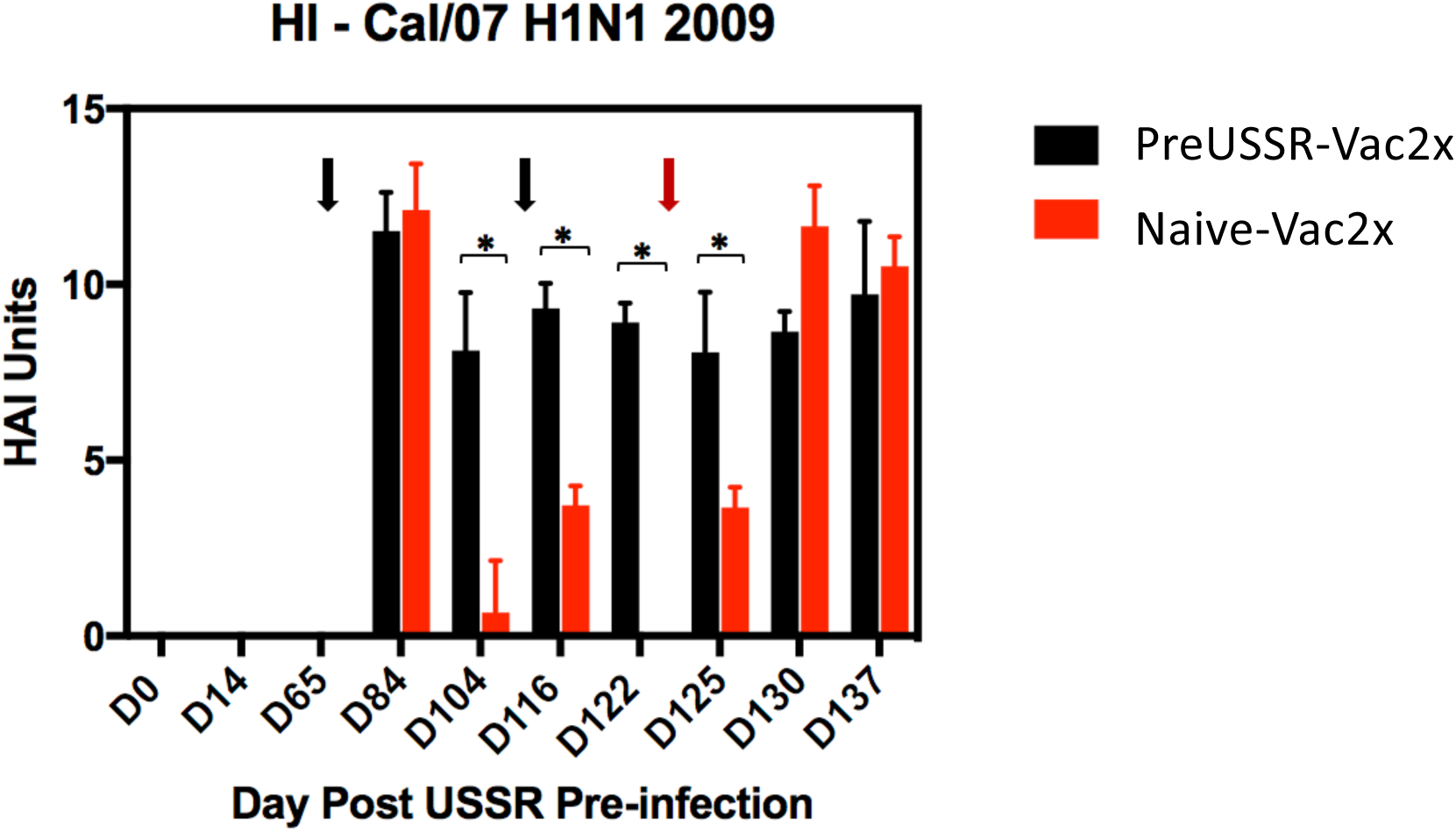
Focused titer analysis of the H1-2009 specific antibody titers shows specific antibody regulation dependent on previous influenza virus exposure. HAI values of only the preUSSR-Vac2x group was compared numerically by histogram to the naïve-Vac2x over the entire time course against the 2009 H1N1 pandemic Cal/09 virus to focus the analysis on the differences caused by immune background/pre-immunity. Black arrows denote time of vaccination. Red arrows denote challenge day. Day 0 is the time of infection with the seasonal H1N1 USSR/77 virus. Values are graphed in HAI Units.

### Mature B cells play a role in the vaccine responses of the imprinted host

Above, we found that the ferrets previously exposed to an antigenically divergent influenza virus had stronger antibody responses to Sanofi QIV^®^ vaccination. From these findings, we next investigated the dynamics of the antibody isotype produced during the sequential infections and vaccinations of our study. To do this, we performed virus specific IgG and IgM ELISAs to quantify the isotype responses specific for the Cal/09 virus. Serum collected from all time points were evaluated but only the time points prior to challenge and following challenge (D2, D7, and D14) are shown (**Figure 7**). No statistical differences were found in the IgM levels among each group throughout the time course with the exception of the naïve-MockVac group did not have any virus specific IgM antibodies until after challenge (left graph) as expected. All IgM titers remained oscillating around 0.5 OD 492 nm. Quantification of virus-specific IgG production found statistical differences in IgG levels among the groups Day 2 and Day 7 pc (right graph). Specifically, the preUSSR-Vac2x group had statistically higher IgG levels (OD =∼1) than all other groups, which had low values close to 0. Naïve-MockVac group were negative for IgG antibody production until after challenge as expected. Interestingly, the preUSSR-MockVac group had significant increases in IgG on Day 7 following infection.

**Figure 7.**
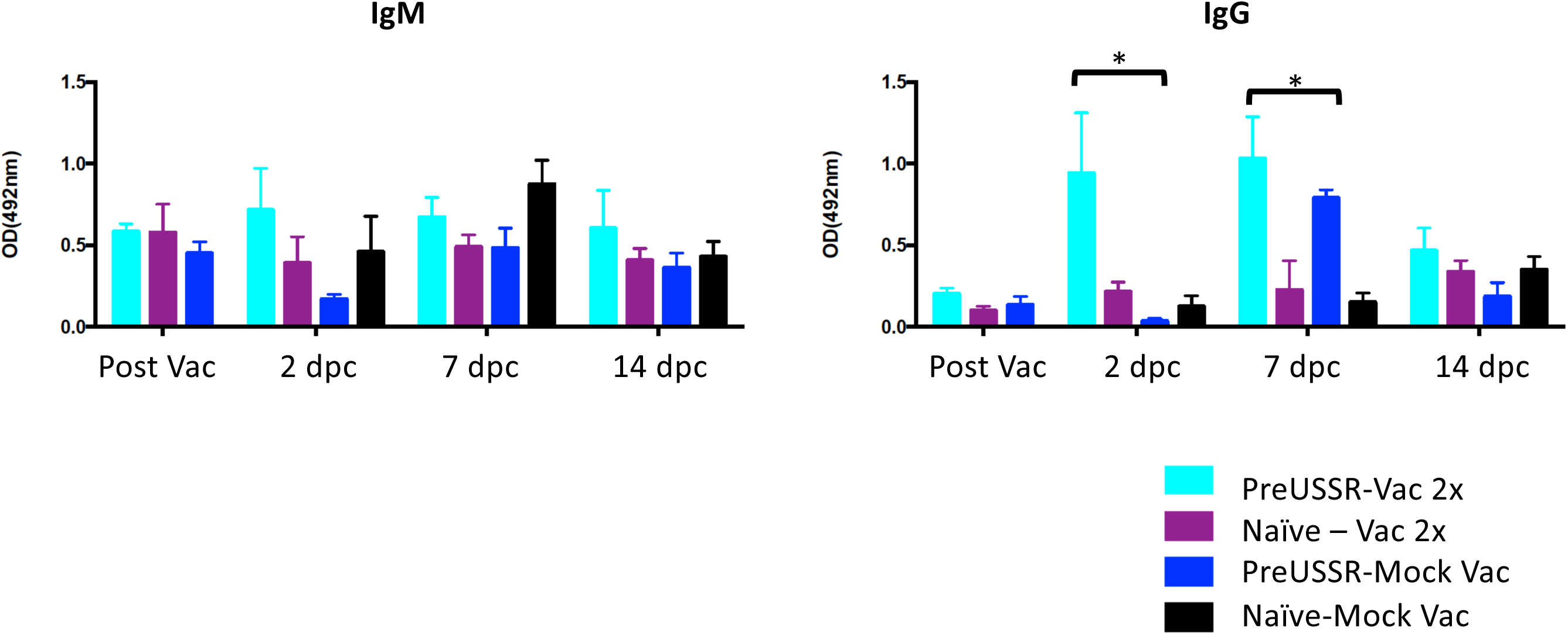
Virus specific IgG isotype antibodies were predominant in the preUSSR-Vac2x group. Isotype ELISAs were performed using the serum collected from each group throughout the time course to determine the proportion of virus-specific IgG or IgM antibodies produced. Only time points post-vaccination and challenge are shown. Student’s t-test was conducted to compare with results on Day 0, \**p*<0.05.

Since the dominant antibody response was of the IgG isotype that is indicative of a more mature B cell response, we were interested in determining the temporal generation of vaccine antigen-specific antibodies at early time points following vaccination. We collected serum in the early time points following vaccination (Days 0, 3, 7, and 14) from naïve ferrets, ferrets previously infected with USSR/77, and naïve-mock vaccinated ferrets as control (**Figure 8A**). HAI assays were performed with the serum against the 2009 H1 antigen and showed that all groups were negative for 2009 H1 HAIs at Days 0 and 3 suggesting that cross-reactive HAI antibodies to the 2009 H1 HA head were not present at these times. By Day 7 the preUSSR-Vac group had a positive HAI response to 2009 H1, reporting at ∼5 HAI Units (light blue bars), whereas both the vaccinated alone and mock vaccinated group did not have a positive HAI. Furthermore, by Day 14 the preUSSR-Vac group had a strong HAI response at 9 HAI Units compared to the naïve-vaccinated ferrets which had a mean of 3 HAI Units for the group (light purple bars). These results suggest that the ferrets with a previous but antigenically divergent H1N1 infection generated a faster and stronger response to the new 2009 H1 antigen.

**Figure 8.**
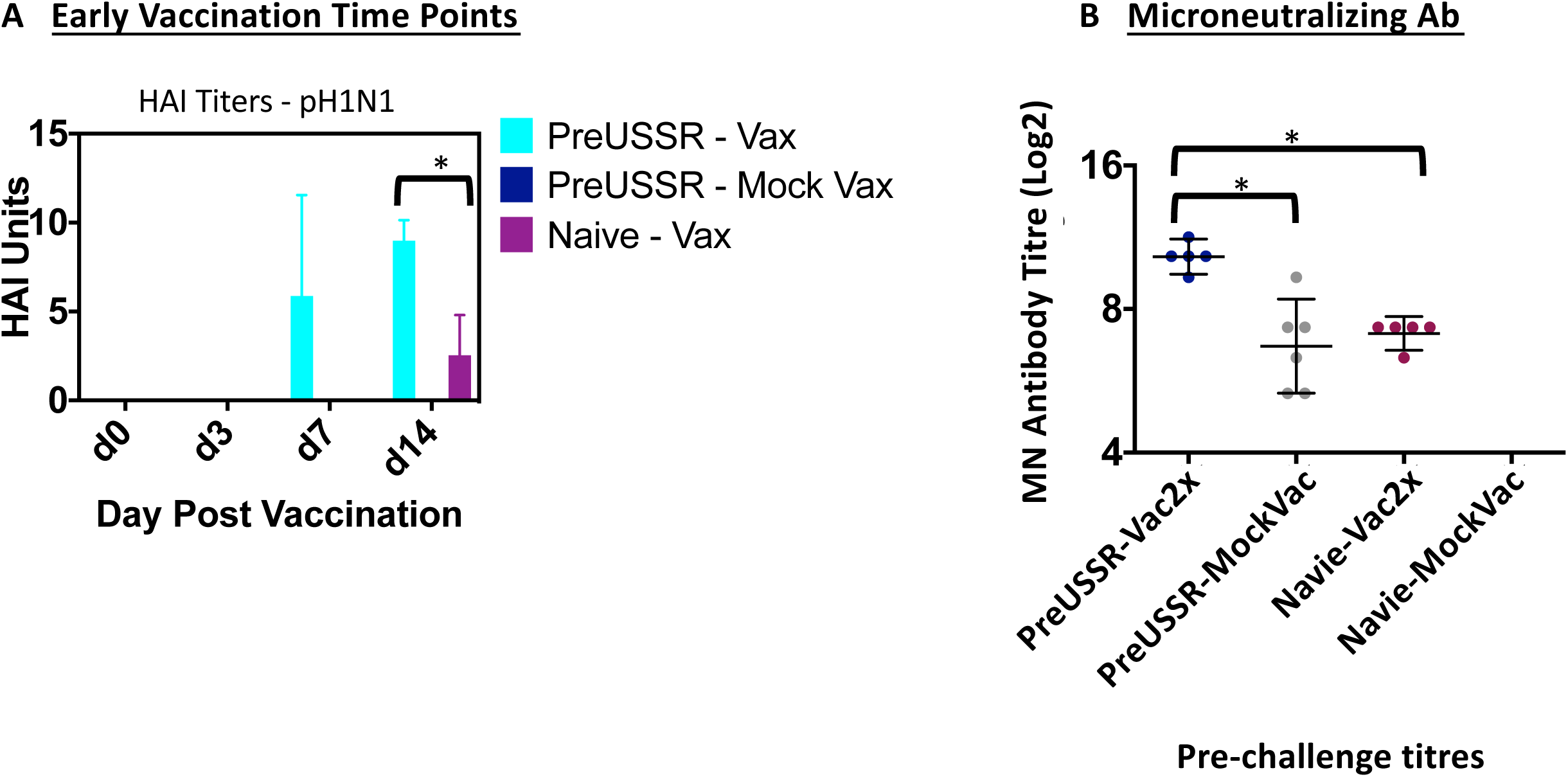
PreUSSR-Vac2x ferrets had specific antibody function and dynamics following vaccination. Serum was collected at early time points post vaccination (Days 0, 3, 7, and 14 post vaccination). HAI assays were performed as previously described on serum against the H1 antigen to determine the early dynamics of the antibody response after vaccination (A). Microneutralization assays were performed by standard protocol on MDCK cells using serum collected prechallenge. Student’s t-test was conducted to compare with results on Day 0, \**p*<0.05.

To investigate the possible mechanism leading to greater protection in the preimmune-vaccinated ferrets, we next investigated the function of the antibodies produced over the time course by performing microneutralization (MN) assays (**Figure 8B**). Using serum collected pre-challenge, MN were performed as previously described against the Cal/09 virus [29]. Pre-challenge serum (Day 123) was used for the MN assays as an indicator or predictor of clinical disease observed during challenge. The preUSSR-Vac2x group had the highest titer of MN antibodies at ∼10, which were statistically greater than any other experimental group. Furthermore, both the preUSSR-MockVac and the naïve-Vac2x had lower MN titers compared to the preUSSR-Vac2 group but similar MN titers to each other. As expected, there were not any MN titers detected for the naïve-mock group. Together, these data show that the preUSSR-Vac2x group had the highest titer of functional antibodies capable of inhibiting viral infection.

Previously, sequential infections with divergent influenza viruses leads to the elicitation of the antibodies that are more broadly reactive toward a larger spectrum of antigenically distinct influenza strains [31–33]. Although the induction of broadly neutralizing antibody generation was shown to be through an HA stem mechanism, we were interested in knowing whether the sequential exposure to these viruses had an effect on the specificity and cross-reactivity of the antibodies produced. Therefore, we performed additional HAIs using the antigenically distinct influenza strain A/Taiwan/1/1986 (Taiwan/86) with our serum samples. Taiwan/86 is antigenically removed from both USSR/77 and Cal/09 and has 90% homology to USSR/77 and 62% to Cal/09 as analyzed by the HA receptor binding domain area (a.a. 135-295)[26]. Previously we showed that direct infection with Taiwan/86 does not elicit cross-reactive antibodies toward either USSR/77 or Cal/09 viruses [26]. In this study we also showed that the reverse was true as well, infection with USSR/77 or Cal/09 did not elicit cross-reactive antibodies toward Taiwan/86 as determined by HAI. These results suggest that any cross-reactivity observed in the sequential infection study would be elicited from repeated influenza antigen exposure. An HAI assay was performed using serum collected at Day 14 pc (Day 137 of the entire study) since this sera would be representative of the largest number of sequential exposures: PreUSSR->QIV->QIVBoost->Challenge. The assay was performed against the Taiwan/86 virus and the results showed distinct trends in HAI cross-reactivity related to the sequence and multiplicity of exposures. Specifically, the preUSSR-MockVac ferrets had the highest titer of cross-reactive antibodies against Taiwan/86 at 9 HAI Units (**Figure 9**). Both vaccinated groups had successively lower titers showing an inverse relationship to the number of repeated exposures. Specifically, the preUSSR-Vac1x and preUSSR-Vac2x had average group titers of 7 and 6 HAI Units, respectively. This data suggested that a low number of sequential exposures could elicit a high titer of cross-reacting antibodies to a highly divergent strain but the more influenza exposures that occur afterward will refine the response.

**Figure 9.**
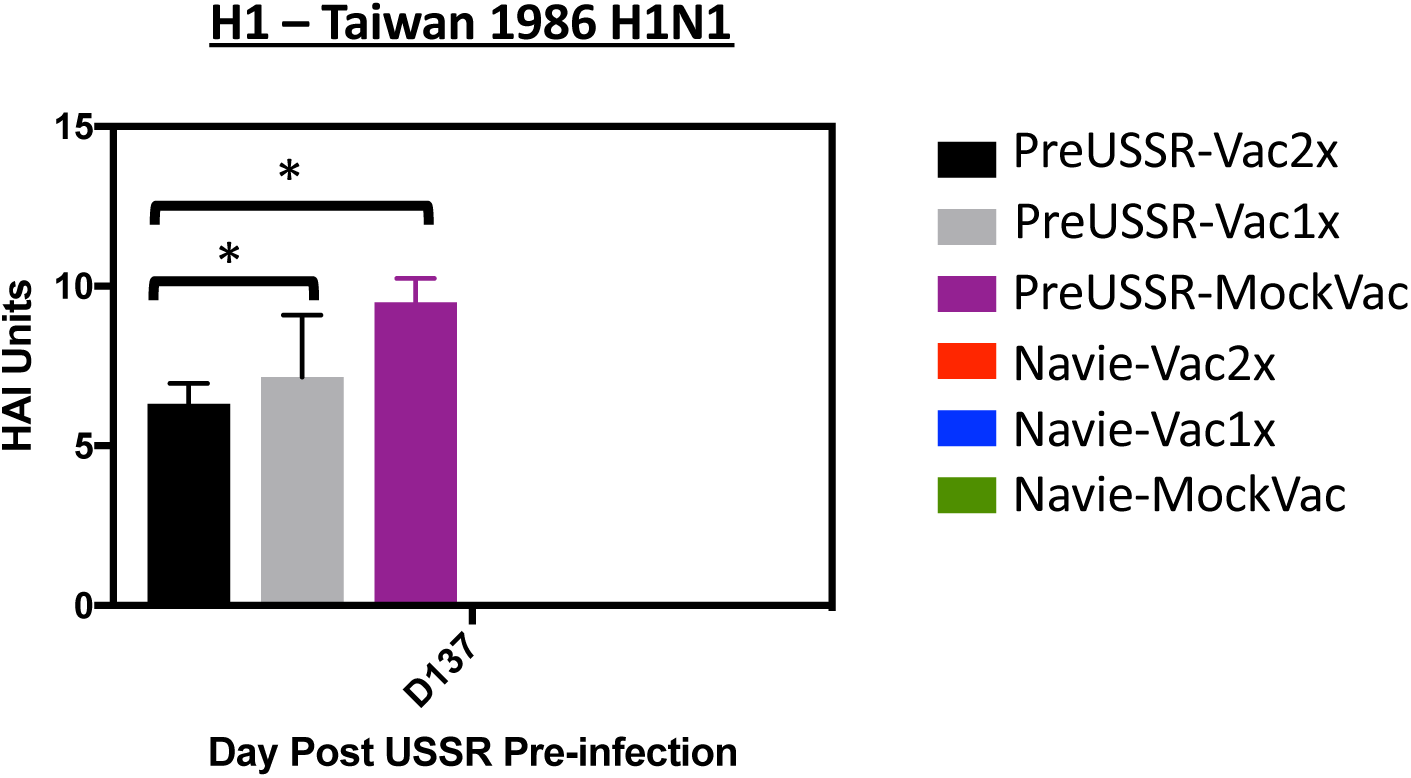
Decrease in cross-reactive antibody titer over sequential influenza virus exposures of antigenically divergent strains. Serum was collected at end day (Day 137) from all ferrets. Serum was then used for HAI assays against an antigenically divergent influenza virus strain (A/Taiwan/1/1986, Taiwan/86) as a read-out for the development of cross-reactive antibodies. Standard HAI assays were conducted with turkey RBCs against Taiwan/86. Student’s t-test was conducted to compare results with the PreUSSR-Mock Vac group, \**p*<0.05.

## DISCUSSION

Every year the influenza vaccine has variable effectiveness against the seasonal circulating viruses allowing continual circulation in humans. Due to continual circulation and influenza virus exposure, humans have a complicated influenza background. The split viron vaccine is the vaccine most often used for seasonal influenza vaccination worldwide, but there has been limited analytical and experimental evaluation in animal models that have been previously exposed to influenza viruses and exhibit with specific influenza memory. To investigate the host-responses to the Sanofi QIV^®^ vaccine, we developed a ferret model with the USSR/77 influenza virus background for vaccine investigation in the immune memory phase of the imprinting infection. We found that the pre-exposed ferrets had increased antibody responses to vaccination that led to increased protection from a divergent challenge virus. Interestingly, although the naïve ferrets were capable of eliciting an antibody response to the vaccine, they developed significant disease with higher viral titers. Together, our results suggest that much could be learned from the immune responses to vaccination in the previously infected host and that the mechanisms identified could be leveraged for improving vaccine responses.

Here our results suggest that previously infected ferrets are better immunologically equipped to respond to the split virion influenza vaccine. We showed that the ferrets preimmune to the USSR/77 virus were able to mount a greater and longer sustained antibody response as determined by HAI assay toward all vaccine antigens. Furthermore, the antibodies elicited had greater neutralization activity suggesting that the mechanism of milder disease at challenge was the ability to develop more functional and viral infection inhibiting antibodies. This is an important result, especially since the naïve-vaccinated ferrets developed a more severe disease compared to other groups. Furthermore, these results suggest that vaccination on a naïve background may lead to the development of antibodies that are not as efficient for inhibiting viral infection. Infants are the most immunologically naïve population in respect to previous influenza virus exposures. Infants also have been shown countless numbers of times to be at a greater risk for developing severe complications requiring hospitalization following influenza virus infection [9]. The higher attack rates in this age group is thought to be partially due to the decreased protection offered from vaccination and it is speculated that the influenza naïve infant immune system is the cause of poor vaccine responses [34]. Our data indicating decreased protection following vaccination in naïve ferrets reflects the ineffectiveness of influenza vaccination in naïve infants suggesting the host may need to be primed prior to vaccination. Since the split virion vaccine was not able to elicit a high level of neutralizing antibodies in naïve ferrets as what was seen in the preimmune ferrets, this suggests that the digested viral antigens of the vaccine is not the most appropriate for inducing functional antibodies. Understanding how a prior infection primes the immune system for more effective vaccinations may offer new strategies for priming the immune system in the naïve hosts. More work is needed to elucidate the molecular mechanisms of antigen elicitation of neutralization antibodies for the development of more effective vaccine platforms.

Our findings also showed that there was an increased disease associated with vaccination on a naïve background. Taken together with our conclusions from the previously exposed and vaccinated group, this data suggested that unprimed individuals may be improperly imprinted during vaccination. Improper viral imprinting by vaccination has also been observed for the respiratory syncytial virus vaccine but at a greater clinical cost [35, 36]. In the 1960s, a formalin-inactivated RSV vaccine was developed and administered to young children. Unfortunately, the formalin inactivation step inhibited subsequent cytoplasmic processing of the viral antigens upon vaccination. The consequence of improper vaccine imprinting was that the vaccinated naïve children developed an ineffective immune response and subsequent infection with RSV led to enhanced disease. Specifically, enhanced disease was dominated by a Th2 response and the vaccine-elicited antibodies were inefficient at neutralizing the virus [35]. Our observation that the naïve-vaccinated ferrets had decreased production of virus neutralizing antibodies suggested that antibodies elicited in the naïve hosts following vaccination may be less functional that those elicited in the preimmune-vaccinated host. Further investigation is needed to determine how to properly imprint the host during vaccination to elicit antibodies that are optimal at inhibiting viral infection.

During influenza virus infection, a large and diverse pool of antibodies are initially produced and circulated [13]. The majority of these are directed toward the HA molecule [37, 38]. The initial humoral response to infection involves the generation of short-lived plasma cells (low-affinity antibody producing cells) that reside in secondary lymphoid organs [16]. These cells expand rapidly, then decline prior to memory establishment. In the germinal centre, B cell specificity is refined for improved responses toward the pathogen. Refinement occurs through a process called affinity maturation. Affinity maturation includes somatic hypermutation (SHM), class-switch recombination (CSR), and clonal selection [39]. Our results from the antibody isotype ELISAs and early time point HAIs showed that preimmune ferrets had a quicker development of vaccine antigen specific antibodies following vaccination and a predominant phenotype of the IgG isotype. The dominance of the IgG isotype suggested the presence of a pre-existing B cell with either existing vaccine antigen specificity or the presence of a pre-existing B cell with the flexibility to modify antigen specificity. Further investigation of the affinity maturation process during sequential exposure of antigenically diverse antigens should be approached to understand how much of a role affinity maturation plays in continual influenza virus infection and vaccination. On the other hand, outside of B and T cell specific responses, there are other immune mechanisms that may be responsible for the faster generation of antibodies following vaccination in the preimmune host as we observed. Specifically, trained immunity is the adaptation of innate immune components after pathogen stimulation. Trained immunity mainly involves the modification of cells such as NK cells, monocytes, and macrophages to produce a “memory” response which can be mobilized over a longer period of time for a second encounter with a specific pathogen [40]. It is possible that either trained immunity or memory B cell dependent responses influenced the specificity of the elicited antibodies after vaccination in the preimmune hosts; however, considering our long rest period between infection and vaccination, trained immunity is less likely. Further investigation of mechanisms of B cell memory and refinement compared to trained immunity will be important to determine the immune mechanisms. Understanding the roles of both mechanisms of memory B cells and trained immunity in the preimmune host’s responses to vaccination may be critical for developing the next generation of influenza vaccines.

Due to the nature of the drifting and shifting influenza viruses, there has been much work investigating immune responses in infection-reinfection animal models. Since the emergence of the pandemic 2009 influenza virus, there have been several experimental and human investigations on monosubtypic heterologous infection-reinfection due to the occurrence of this infection sequence in the human population. In summary, the general finding from these studies was that primary infection with seasonal H1N1 from the 1977 H1N1 lineage followed by an infection with pandemic 2009 virus leads to partial protection with reduction in disease clinical severity in infected individuals [41–46]. We previously reported that primary infection induced pre-existing non-HA antibodies during 2009 H1N1 virus challenge due to sequence homology with the less immunogenic internal viral proteins. Other reports showed this sequential infection leads to the generation of both HA head and stalk directed antibodies and that stalk antibodies may be responsible for a more broadly protective antibody response [31–33, 47]. Together, these reports suggest that antibodies generated toward internal proteins and/or the HA stem may also play a role in the clinical outcome we observed. Although we can glean information on the immune responses in the pre-exposed hosts from the experimental infection-reinfection studies, it is important to note that our study involved sequential viral antigen exposure both in the forms of live viral infection and inactivated viral antigens in the split virion vaccine. Our study specifically showed a broader antibody responses in the preimmune ferrets directed toward the HA head as shown in our HA assays which was not recognized in these other studies. From our data we are the first to show evidence for a HA head directed mechanism of protection which is possibly due to the digested viral antigens of the vaccine. Future investigation of the antibodies elicited toward the HA stem and viral internal proteins using the long recovery model should also be done similarly to the live viral sequential infection studies [32, 33] to gain a more complete picture of the humoral immune responses in the preimmune host during vaccination.

Our study specifically investigated the vaccine responses in hosts that had a previous seasonal H1N1 1977 lineage virus infection and then a 2009 pandemic H1N1 virus challenge following vaccination. Since these viruses are of the same subtype but different lineage, it is unknown how vaccines responses may be regulated on a different influenza virus background. Each influenza season is dominated by a specific circulating subtype or lineage, suggesting that the imprinting exposure event of a person may be equally from an H3N2, B-Yamagata, or B-Victoria virus strain [9]. Antigenically as well as biochemically in terms of glycosylation, these virus subtypes and lineages are very different from the H1N1 virus which may impact the imprinting response [48–50]. Future studies should investigate how the host imprints on these viruses and how the immune responses to a subsequent vaccination are regulated. Studies such as these will give insight into which human cohorts will respond to particular vaccine formations. This information will be important for identifying susceptible human populations during virulent influenza seasons and pandemic virus emergence.

Although great efforts have been made to develop effective influenza virus vaccines, there has been little progress improving vaccine effectiveness as evidenced by the moderate overall effectiveness of the 2018-2019 vaccine at 47% [51]. Several vaccine formulations now exist but it is not known how the preimmune host will response to these platforms. The Centers for Disease Control and Prevention estimated over 80,000 deaths occurred due to influenza virus infection in the 2017-2018 influenza season. Our study takes a step forward to understanding the actual vaccine responses in the majority of humans, since all people over the age of 4 will have an influenza immune history [14]. Together, our findings suggest that H1N1 influenza immune priming leads to more efficacious vaccine responses. Determining how to prime the immune system without the deleterious effects of infection should now be an essential research goal. Importantly, our results may only be extrapolated to human situations with a single previous infection prior to vaccination. Using our strategy, further work is needed to tease out the efficacy and vaccine responses on alternate influenza virus backgrounds such as H3N2 or a layered serial influenza background where sequential infections lead to immunesenescence [52]. As well the responses to vaccination with other vaccine platforms such as the Live Attenuated Influenza Virus (LAIV) vaccine should also be investigated.

## METHODS

### Ethics statement

All animal work was conducted in strict accordance with the Canadian Council of Animal Care (CCAC) guidelines. The protocol license numbers AUP 5316 and 1031 were assigned by the Animal Care Committee of the University Health Network (UHN). UHN has certification with the Animals for Research Act, including for the Ontario Ministry of Agriculture, Food and Rural Affairs, Permit Numbers: #0132–01 and #0132–05, and follows NIH guidelines (OLAW #A5408-01). The animal use protocol was approved by the UHN Animal Care Committee (ACC). All efforts were made to minimize animal suffering. Infections and sample collections were performed under 5% isoflurane anesthesia.

### Influenza Virus and animals

The 2009 H1N1 virus strains, A/California/07/2009 (Cal/07), A/USSR/90/1977 (USSR/77), and A/Taiwan/1/1986 (Taiwan/86), were provided by the Influenza Reagent Resource, Influenza Division, WHO Collaborating Center for Surveillance, Epidemiology and Control of Influenza, Centers for Disease Control and Prevention, Atlanta, GA, USA. TCID_50_ and EID_50_ determinations were done as previously described [53]. All virus work was performed in a BSL-2+ facility as previously described [29]. Sanofi FLUZONE^®^ quadrivalent QIV influenza vaccine from the 2015-2016 influenza season was acquired from Sanofi Canada (North York, Canada). The vaccine contained concentrated HA proteins from A/California/07/2009 (H1N1); A/Victoria/210/2009 (H3N2); B/Brisbane/60/2008 (B-Victoria Lineage); B/Florida/04/2006 (B-Yamagata Lineage). Adult female ferrets (aged ∼5 months to 1 year) were purchased from Triple F Farms (Gillett, PA, USA). Ferrets were determined to be seronegative by haemagglutination inhibition (HI) assay against currently circulating influenza A and B strains before infection.

### Infections and Vaccinations

Ferret were infected intranasally as previously done [54]. Briefly, ferrets were anesthetized and infected with seasonal viruses or pandemic viruses at 10^6^ EID_50_. The volume of inoculum was 1 mL for each ferret (0.5 mL in each nare). For vaccination, ferrets were vaccinated intramuscularly with a whole human dose of FLUZONE^®^ Sanofi QIV vaccine in the upper hind limb. Ferrets were observed post infection or vaccination for adverse effects.

### Clinical Monitoring

Weight, temperature, and clinical signs were monitored following infections and vaccinations for 14 days similarly as previously done [55]. Weight and temperature for each day were calculated as a percentage of original values determined on Day 0 and days prior to study initiation. Standard deviation and standard error were calculated for weight and temperature percentages within each group. Clinical signs (body temperature, body weight, level of activity, nasal discharge, and sneezing) were observed daily for 14 days pi, post-vaccination, or pc. Ferrets were examined at the same time each day for consistency. Nasal discharge, sneezing, and inactivity was observed and recorded but not reported here.

### Viral Titers, HAI Titers, Microneutralization titers (MN), and IgG/IgM relative isotype levels

Viral titers were calculated from collected nasal wash samples pc. Nasal washes were subjected to a TCID50 assay followed by HA assay against virus strain of interest to determine viral titers. Viral load was calculated using the Reid and Meunch method [30]. HAI titers were determined in serum collected throughout the entire study by HAI assay using USSR/77, Cal/09, or Taiwan/86 live viruses or using the WHO circulating influenza virus detection kit (2016-2017). The WHO kit assays for H1N1 2009, circulating H3N2, circulating B-Yamagata, and circulating B-Victoria strains. Microneutralization titers (MN) results were evaluated by enzyme-linked immunosorbent assay (ELISA). Neutralizing antibody titers were determined by the highest dilution of RDE-treated anti-sera that disrupted infection (100 TCID_50_) on MDCK cells at the reading lower than 50% signal reading measured from virus+cell and cell only controls. The determination of IgG and IgM specific antibodies in collected serum was based on enzyme-linked immunosorbent assay (ELISA) technique as previously described [28]. Briefly, ELISA plates were directly coated with A/California/07/2009 overnight at room temperature. Plates were washed with PBS containing 0.05% Tween 20 (T-PBS) and blocked with 1% bovine serum albumin (BSA) for 1 h at 37°C. Antigen-coated plates were incubated with 1:1,000-diluted serum samples overnight at 4°C. After washing with T-PBS, plates were incubated with goat anti-ferret immunoglobulin (IgM and IgG) horseradish peroxidase (HRP) conjugates (Rockland Immunochemicals) in a 1:10,000 dilution for 2 h at 37°C. The reaction was developed by *o*-phenylenediamine for 30 min, and the optical density was read at 492 nm.

### Histopathology

Lungs were collected at necropsy, perfused with formalin and paraffin embedded. Following sectioning the tissues were mounted and H&E stained. High resolution scans were performed using an Aperio ScanScope XT (Leica Biosystems) at 40x magnification. Images were captured using the HALO program from UHN AOMF (Advanced Optical Microscopy Facility) at 5x, 10x, or 20x magnification of the scan.

### Statistical Analysis

ANOVAS or Student’s t-test was used to determine p-values when comparing groups. The inactivity index of each viral infection was calculated using the scores observed daily. For both the viral and antibody titers, standard deviation was calculated within each experimental group. Student’s t-test or ANOVAS was used to determine the p-value among groups.

## ACKNOWLEDGEMENTS

This work was supported by research is funded by the National Institutes of Health NIH 1U01AI11598-01 Subaward no. F8802-16 S and National Institutes of Health NIH Contract HHSN272201000031I, Order Number HHSN27200003, Task A88 AAK and TMR. https://www.nih.gov/ The funders had no role in study design, data collection and analysis, decision to publish, or preparation of the manuscript. Thank you to Luoling Xu, Amber Farooqui, and Adnan Khan for their technical assistance on this project.

## AUTHOR CONTRIBUTIONS

Conceived and designed the experiments: AAK. Performed the experiments: AAK LX AK AF MF MK. Analyzed the data: AAK TMR. Wrote the paper: AAK.

## SUPPORTING INFORMATION CAPTIONS

**Suppl. Figure 1.**
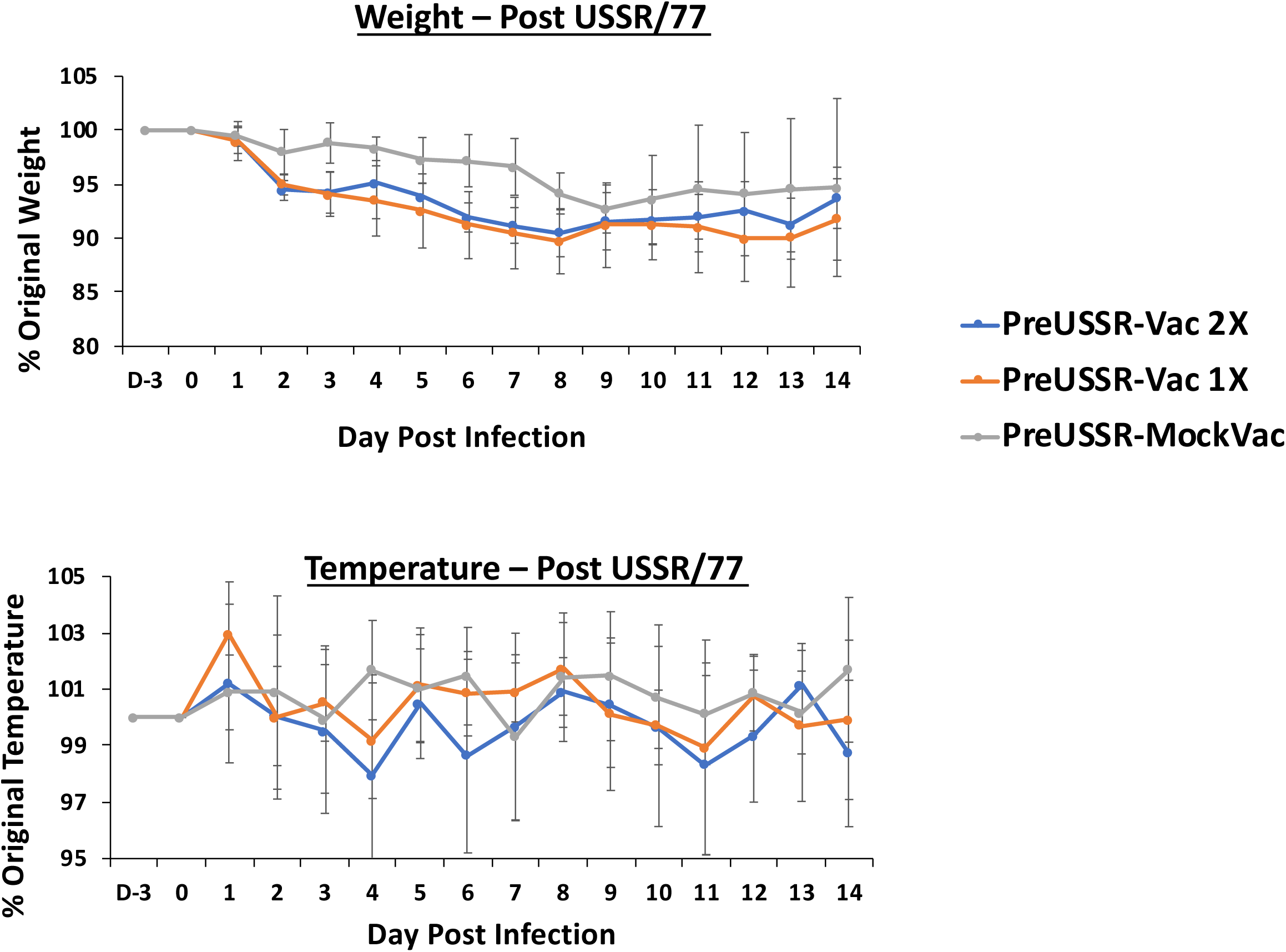
Weight loss and temperature change following infection with A/USSR/90/1977 to establish a preimmune background. Ferrets were intranasally inoculated with A/USSR/90/1977 in PBS at a dose of 10^6^ EID_50_. Ferrets were weighed and temperature was taken daily. The percentage of original values was calculated throughout 14 days.

**Suppl. Figure 2.**
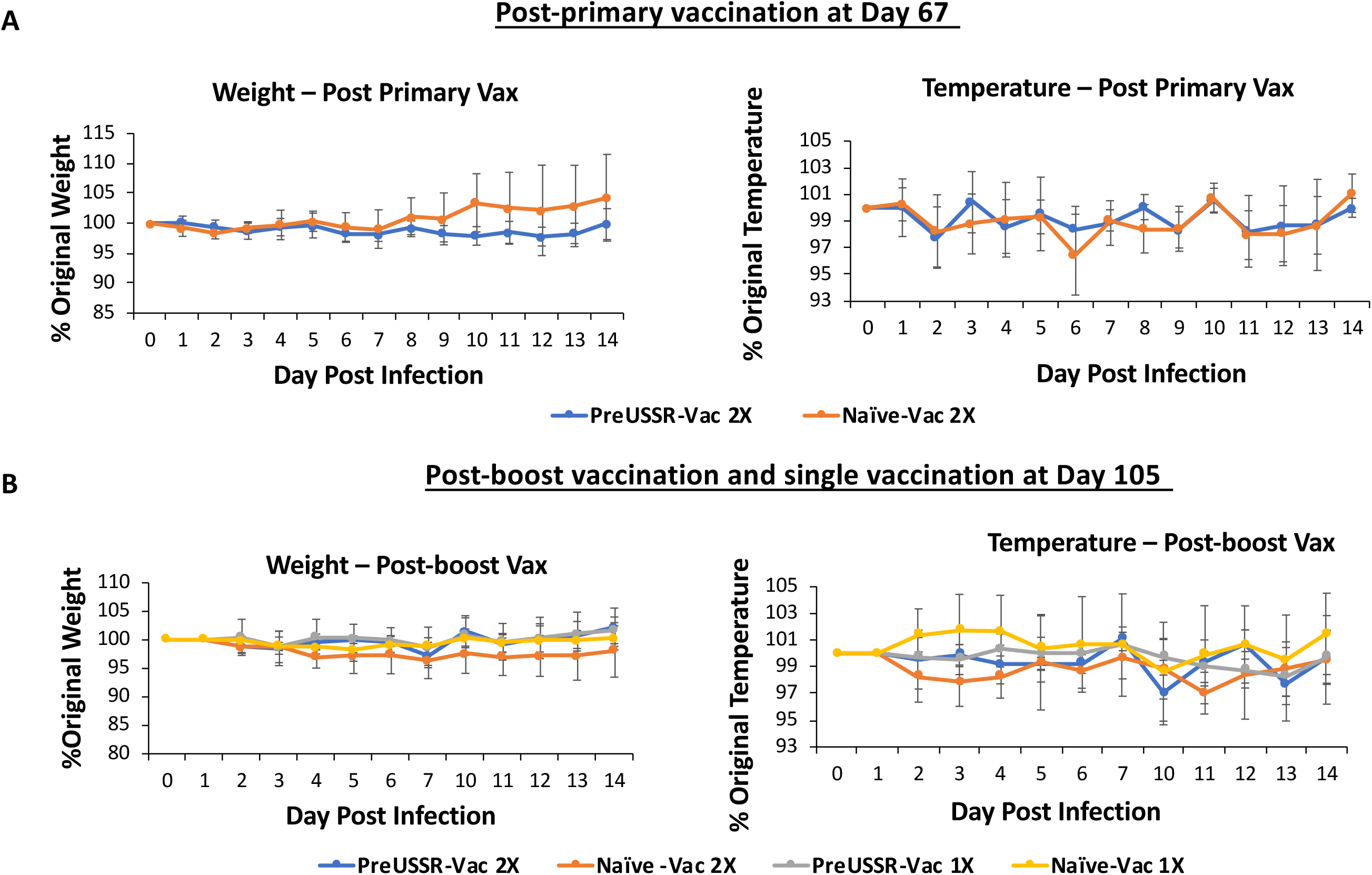
Minimal weight loss and temperature change were observed following vaccination with Sanofi FLUZONE^®^ QIV. Both Naïve and preimmune ferrets were vaccinated by intramuscular injection with the Sanofi FLUZONE^®^ QIV split virion vaccine on Day 67 and/or Day 105 following the initial imprinting infection. Ferrets were weighed and temperature was recorded daily. The percentage of original values was calculated and graphed throughout 14 days.

**Suppl. Figure 3.**
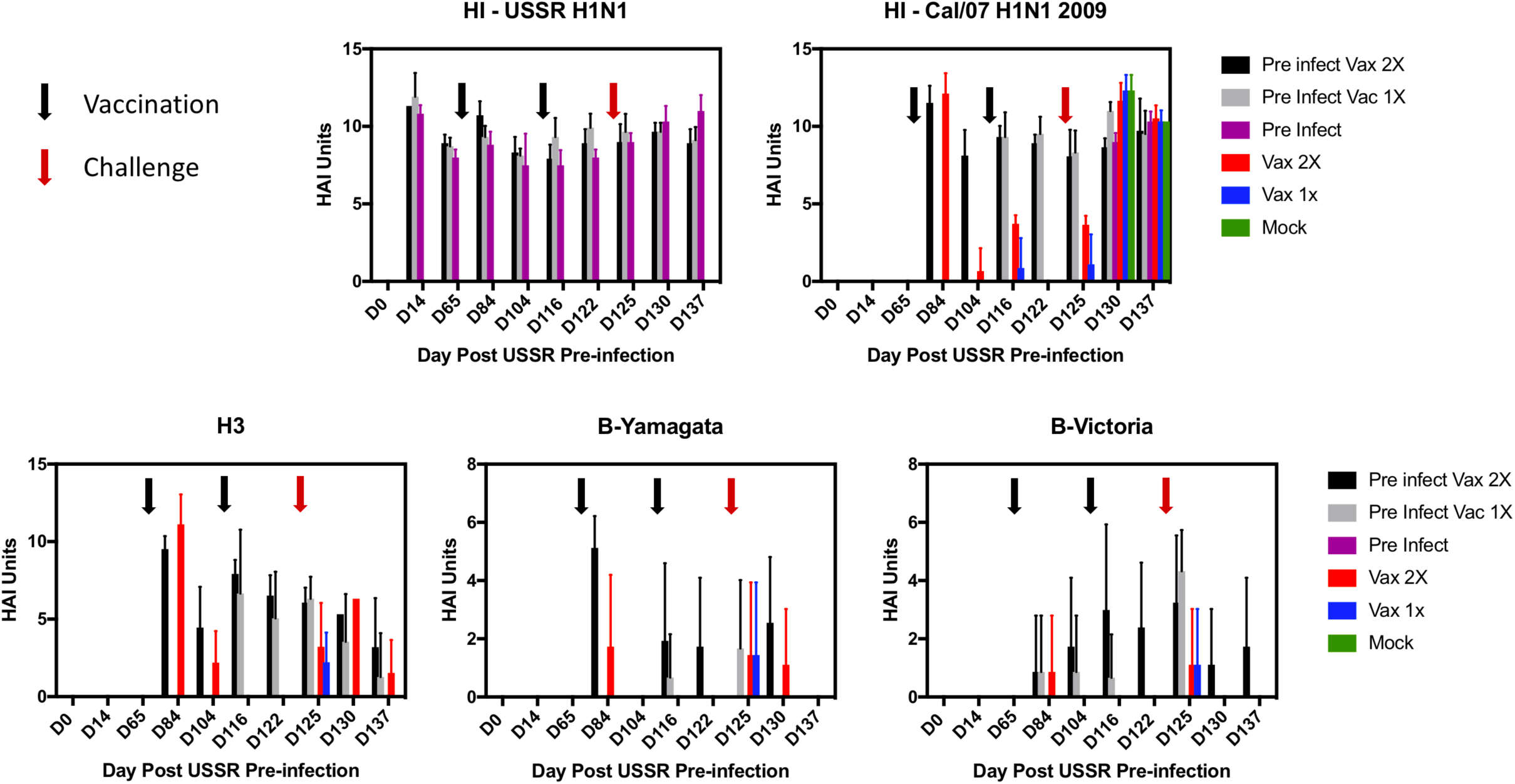
Histogram visualization of HAI titer regulation over the experimental time course showed preimmune ferrets had greater antibody responses compared to naïve-vaccinated ferrets. The HAI U for each experimental group was calculated at each time point for the HAI titer against the Cal/09 virus. The results were plotted as histograms over the infection time course to visualize the numerical dynamics and changes over time. HAI values are plotted per antigen/virus. Time of vaccination is indicated by a black arrow. Time of challenge is indicated by a red arrow. Day 0 denotes the first day of the study (H1N1 USSR/77 virus inoculation day in appropriate groups).

